# Human Genes Escaping X-inactivation Revealed by Single Cell Expression Data

**DOI:** 10.1101/486084

**Authors:** Kerem Wainer Katsir, Michal Linial

## Abstract

**Background:** In mammals, sex chromosomes pose an inherent imbalance of gene expression between sexes. In each female somatic cell, random inactivation of one of the X-chromosomes restores this balance. While most genes from the inactivated X-chromosome are silenced, 15-25% are known to escape X-inactivation (termed escapees). The expression levels of these genes are attributed to sex-dependent phenotypic variability.

**Results:** We used single-cell RNA-Seq to detect escapees in somatic cells. As only one X-chromosome is inactivated in each cell, the origin of expression from the active or inactive chromosome can be determined from the variation of sequenced RNAs. We analyzed primary, healthy fibroblasts (n=104), and clonal lymphoblasts with sequenced parental genomes (n=25) by measuring the degree of allelic-specific expression (ASE) from heterozygous sites. We identified 24 and 49 candidate escapees, at varying degree of confidence, from the fibroblast and lymphoblast transcriptomes, respectively. We critically test the validity of escapee annotations by comparing our findings with a large collection of independent studies. We find that most genes (66%) from the unified set were previously reported as escapees. Furthermore, out of the overlooked escapees, 11 are long noncoding RNA (lncRNAs).

**Conclusions:** X-chromosome inactivation and escaping from it are robust, permanent phenomena that are best studies at a single-cell resolution. The cumulative information from individual cells increases the potential of identifying escapees. Moreover, despite the use of a limited number of cells, clonal cells (i.e., same X-chromosomes are coordinately inhibited) with genomic phasing are valuable for detecting escapees at high confidence. Generalizing the method to uncharacterized genomic loci resulted in lncRNAs escapees which account for 20% of the listed candidates. By confirming genes as escapees and propose others as candidates from two different cell types, we contribute to the cumulative knowledge and reliability of human escapees.

## Background

Sex chromosomes pose an inherent genetic imbalance of gene expression between sexes. In order to ensure a balanced expression in mammalian somatic tissues, one of the female’s X-chromosomes (ChrX) is randomly selected to undergo inactivation [1]. The random choice of an inactivated X-chromosome (Xi) (*i.e*., paternal or maternal) is completed at a very early phase of embryonic development [2]. Importantly, once this decision is made the selected inactivated chromosome is deterministically defined for all descendant cells, and this choice is maintained throughout the organism’s life in every somatic tissue [3]. This highly regulated process has been extensively studied [2-5].

The initial silencing of ChrX is governed mainly by *XIST* (X-inactive specific transcript) [3, 4], a non-coding RNA (ncRNA) unique to placental mammals. *XIST* is a master regulator located at the X-inactivation center (XIC) that together with neighboring ncRNAs (e.g., *FTX* and *JPX*) activate the process of X-inactivation [3]. *XIST* is exclusively transcribed from Xi, and its RNA products act in cis by coating the chromosome within a restricted chromosomal territory [6]. The activity of XIC genes in recruiting chromatin remodeling complexes [3, 7, 8], results in an irreversible heterochromatinization. The heterochromatin state underlies the steady, lifelong phenomenon of X-inactivation [1].

Ample studies have indicated that silencing does not apply to all genes in the inactivated X-chromosome. Specifically, genes that are located at the Pseudoautosomal regions (PARs) are expressed from both alleles, similar to the majority of genes from autosomal chromosomes [9]. In addition, on the ChrX there are also genes that escape X-inactivation (coined escapees). Investigating these escapee genes is important to understand the basis of ChrX evolution [10] and X-inactivation mechanism [7]. Moreover, numerous clinical and phenotypic outcomes are thought to be explained by the status of escapee genes [11].

Complementary methods have been adapted for identifying escapees [12, 13]. For example, the expression levels of mRNAs were compared between males and females in various tissues [14-16]. Additionally, extensive lists of escapee candidates were reported from mouse-human cell hybrids, and from allelic expression patterns in fibroblast lines carrying a fragmented X-chromosome [17]. The correlation of chromatin structure and CpG methylation patterns with genes that escape X-inactivation was also used. For example, loci on Xi with low methylation levels were proposed as indicators for escapee genes and were thus used as an additional detection method [18, 19].

In recent studies, genomic information from individuals and isolated cells became useful for marking the status of X-inactivation. Specifically, RNA sequencing (RNA-Seq) was used to infer allelic-specific expression (ASE) from the two X-chromosomes, according to a statistical assumption for the minor and major expressed alleles [20]. ASE analysis from B-lymphocytes derived from two ethnically remote populations identified 114 escapees based on heterologous SNPs (hSNPs) [10]. By default, the low-expressing hSNP alleles were considered as evidence for Xi expression. Recently, a large-scale ASE-based analysis was completed based on a few individuals using single cells [16].

Numerous observations indicate conflicts and inconclusive labeling of a ChrX gene as inactivated or escapee. Such variability reflects the inherent properties of the phenomenon with respect to tissues, individuals and developmental stages. Several trends characterize X-inactivation and escaping from it: (i) Escapees are located at the p-arm, which comprises evolutionary young segments that diverged more recently from ChrY [17, 21, 22]. (ii) Human escapees account for 15-25% [13]) of all known ChrX genes. Notably, this fraction is only ~3% for mice [11, 23]. (iii) A low level of expression signifies Xi when compared to Xa (e.g., [10, 16, 24]). (iv) Most escapees show a substantial heterogeneity across cell types, individuals and experimental settings [13, 16, 24, 25]. (v) Only a few of the escapees exhibit consistent behavior across a wide range of cell types and conditions [16, 24]. (vi) Some clinical and phenotypic diversity result from the varying degree of X-inactivation and escapee’s expression (e.g., [10, 26]).

The X-inactivation is an event that occurs independently for each cell. Thus, collecting expression data from single cells allows monitoring explicitly a single Xi allele in each cell. In the present study, we use the ASE data extracted from RNA-Seq of single cells (scRNA-Seq) for identifying escapees. We present an analytical protocol using genomic data for two sets collected from scRNA-Seq experiments. One set is based on primary fibroblasts, and the other is based on GM12878 lymphoblast cell line with a fully sequenced diploid genome. We report on 24 genes from fibroblasts and 49 genes from lymphoblasts as candidate escapees. Finally, we demonstrate the potential of the method to identify a large number of escapees despite a modest number of single cells analyzed. We show that while most of our identified escapees strongly agree with the current knowledge, we also provide an extended list of escapes that were previously undetected.

## Results and Discussion

### A framework for measuring the escape from X-inactivation in single cells

We identify escapees by analyzing gene expression from somatic single cells using scRNA-Seq methodology (see Methods). To evaluate the sensitivity of the method, we compare X-chromosome (ChrX) expression to other autosomal chromosomes. Specifically, we focused on the gene-rich chromosome 17 (Chr17) as a prototype of an autosomal chromosome. Chr17 was selected as it represents a chromosome with a minimal number of parent-specific imprinted genes [27]. The quantitative properties of ChrX and Chr17 are listed in Fig. 1a.

**Fig. 1.**
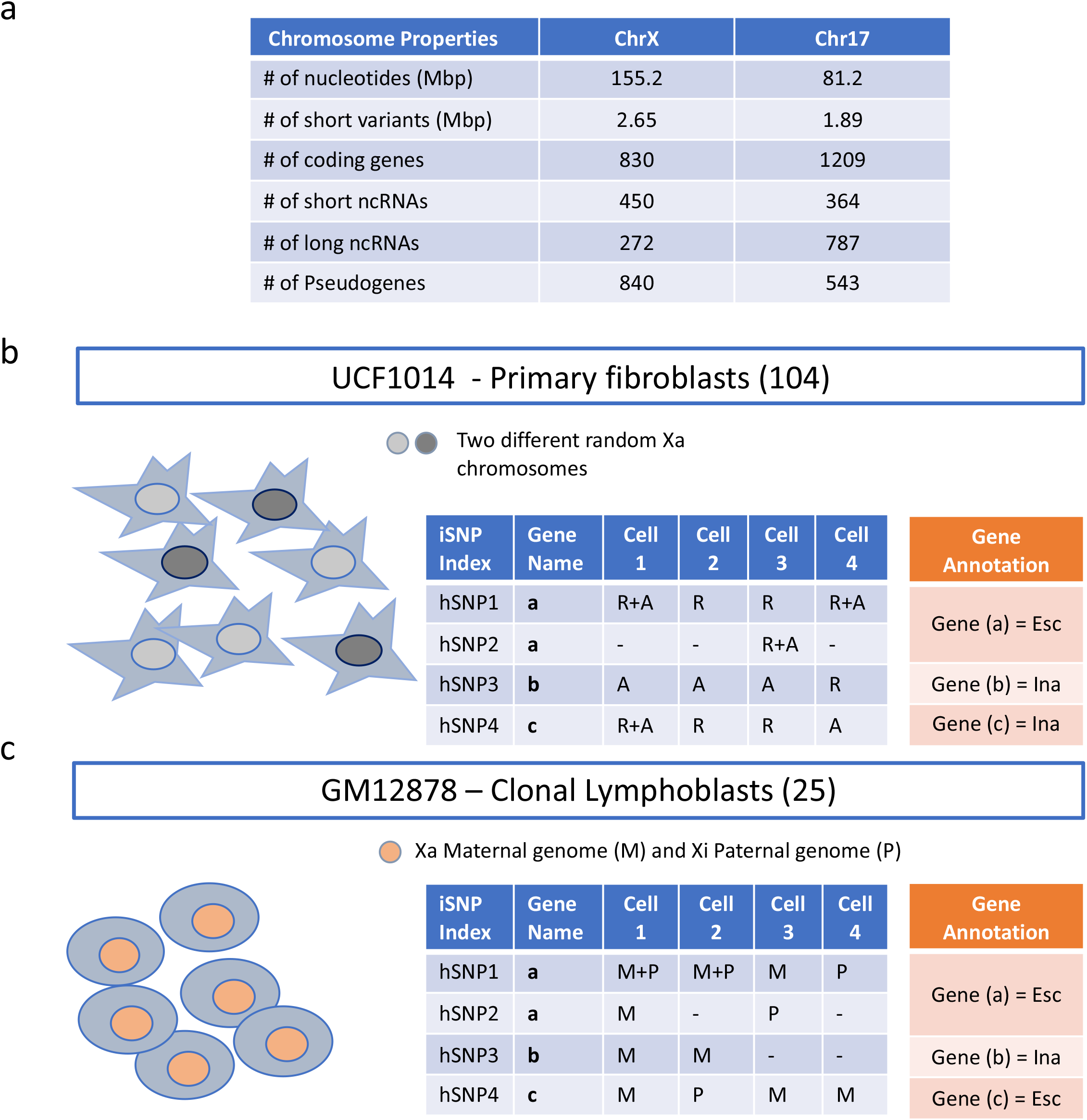
Workflow for identifying escapee genes from single cells. **(a)** Quantitative properties of ChrX and Chr17 are listed according to GRCh37 (GRC Human Build 37). **(b)** A scheme for the analysis of single cell primary fibroblasts. The two colors for the nuclei represent the random choice of Xa. In the context of fibroblasts, each Xa exhibits a different pattern of expression for the hSNPs. Each of the iSNPs can be assigned to the reference (R) or the alternative allele (A). If one cell with one Xa will have an expression pattern of A R A, a cell with the alternative Xa will express R A R. Due to the random X inactivation, and the hSNPs not being phased, annotating a gene as an escapee is entirely based on having multiple evidence of iSNPs with biallelic expression. The illustrative table shows the analysis of iSNPs from each of the hSNPs (on the left) in each of the cells as A or R and the annotation of a gene according to the accumulation of the iSNPs evidence. The illustration marks hSNPs derived from 4 single cells (cell-1 to cell-4). The hSNPs are associated with 3 genes (marked as gene a to gene c). Gene a is the only gene with multiple biallelic iSNPs thus it is annotated as Escapee gene (Esc). The other two genes either do not have biallelic iSNPs (gene b) or have only a single iSNP as evidence for biallelic expression (gene c) and thus are annotated as Inactivated gene (Ina). **(c)** The scheme for the single cells clonal lymphoblasts. In contrast to primary fibroblasts (b), the parental origin of Xa is identical for all cells. In this case of GM12878 cell-line Xa is associated with the maternal (M) allele (symbolized by pink colored nuclei). For lymphoblasts, the occurrence of a paternal allele (marked as P) suffices the identification of an iSNP being expressed from Xi and thus can be potentially annotated an escapee. The table on the right emphasized lymphoblast escapee assignment. The categories of the table are the same as in (b). For details on the workflow and the applied protocols, see Additional file 1: Text and Fig. S1.

This study is based on analysis of two female origin resources: (i) Primary UCF1014 fibroblasts (with 104 cells, see Methods). This set is specified by a higher coverage transcriptomic data, but lacks information on haplotype phasing (Fig. 1b); (ii) A smaller dataset of clonal lymphoblasts (n=25) from the GM12878 cell line with fully phased and sequenced parental diploid genomes (Fig. 1c). In both datasets, transcription at heterozygous SNPs (hSNPs) is the source of information for determining monoallelic or biallelic expression. Each hSNP, in every cell, that is supported by expression evidence above a predetermined threshold is considered an informative SNP (iSNP) (see Methods, Additional file 1: Text). The sum of iSNPs per gene defines its unique label as inactivated or escapee gene (see Methods, Figs 1b-1c, Additional file 1: Text).

### Quantifying biallelic expression from single cell primary fibroblasts

We analyzed the published scRNA-Seq data from female primary human fibroblasts [28]. Before analysis, we took care of an experimental pitfall relevant to many single-cell studies. The pitfall concerns cell doubles in which more than one cell is sequenced per one library. In such a scenario, different active X-chromosomes (Xa) from two different cells that are included in the sample will produce a biallelic signal along the entire X-chromosome. While the fraction of doublets is expected to be small, it may lead to wrong interpretation [29]. Therefore, before analyzing the data from the fibroblasts, we revisited all 104 fibroblasts and tested their biallelic ratio with respect to ChrX (see Methods). Three cells showed an exceptionally high degree of biallelic expression that might indicate a mixture of two parental X-chromosomes (Additional file 1: Text and Fig. S3). We removed all three suspicious cells from all the analyses.

Next, for every single cell, we counted the number of reads that were uniquely mapped to hSNP alleles. The allelic ratio (AR) for each iSNP is defined as the fraction of the reads mapped to the alternative allele (Alt) out of the total reads (see Methods, Additional file 2: Table S1). Figs 2a-2c summarizes the AR of ChrX, Chr17, and the entire autosomal chromosomes according to the primary fibroblasts collection (101 out of 104 cells). In addition, Fig 2d shows the distribution of the AR of an annotated set of imprinted genes from skin tissues (according to [27]). As previously reported [30, 31], a bias in mapping towards the reference genome (AR=0) is evident (Figs. 2a-2d). Additionally, a substantial fraction of monoallelic expression was observed for all tested sets (Fig. 2a-2d). This dominant appearance of monoallelic expression in single cells is caused by a combination of both under-sampling of transcripts, and a phenomenon that is known as “transcriptional bursting” or “dynamic random monoallelic expression” [28, 32-34]. Such transcription bursts reflect the stochastic nature of gene expression in single cells [35].

**Fig. 2.**
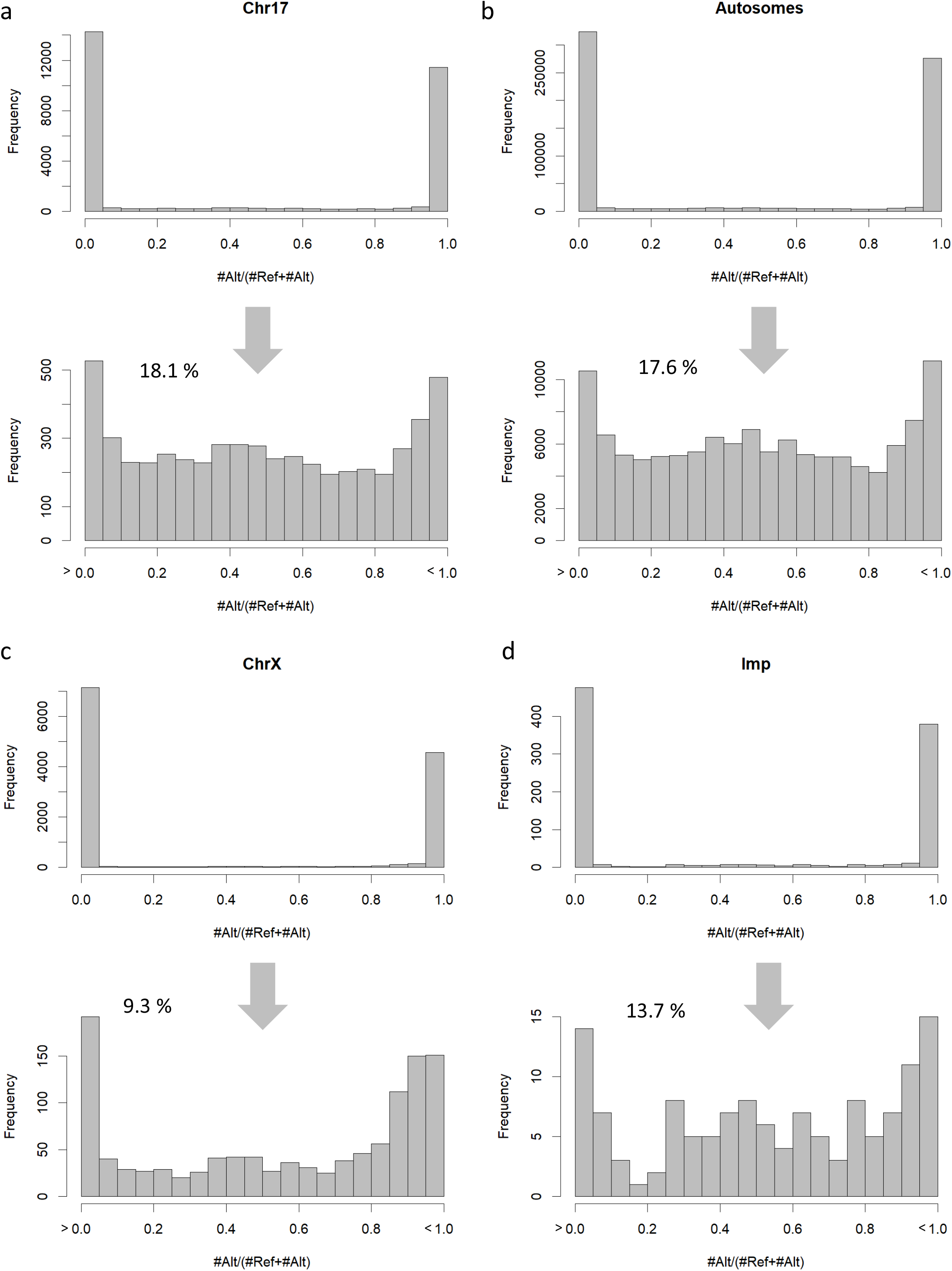
The distribution of the allelic ratio (AR) for each SNP as a fraction of the assignments for Alternative (Alt) out of Alt and Reference (Ref) alleles. The X-axis in the top histograms ranges from 0 to 1.0, where 0 indicates that all assignments are associated with the Ref allele and 1 indicates all assignments for the Alt allele. As the majority of the iSNPs are assigned with AR values of 0 or 1, each analysis is shown by two histograms. The lower histogram focuses on non-monoallelic iSNPs and covers all AR values excluding the AR=0 and AR=1. The percentage of iSNPs that are included in the lower histograms are shown. The distributions of the AR are shown for Chr17 **(a)**, Autosomal chromosomes **(b)**, ChrX **(c)** and imprinted genes **(d)**. For source data, see Additional file 3: Table S2.

We focused only on iSNPs that show a non-monoallelic signature (i.e., excluding AR=0 and AR=1). We observed a marked difference in the AR distribution of ChrX and imprinted genes relative to Chr17 and all autosomal chromosomes (compare Figs 2a to 2b and Fig. 2c to 2d). Accordingly, several observations from the results shown in Fig. 2 can be drawn: (i) Chr17 and all autosomes share a similar AR profile. (ii) A clear tendency towards balanced expression (AR=0.5) is apparent for any autosomal chromosomes (Fig. 2a-2b), but not ChrX or imprinted genes (Fig2 2c-2d). (iii) The fraction of non-monoallelic expression in autosomal chromosomes is higher (~18%) relative to ChrX (~9%). (iv) The fraction of non-monoallelic expression in imprinted genes shows an intermediate level (13%). Such an intermediate level is probably a reflection of the inherent inconsistency in the identity of imprinted genes [36, 37]. We conclude that the attenuation of the non-monoallelic expression for ChrX is a strong indicator of the robustness of the inactivation phenomenon in primary cells. It is worth noting that the strong inactivation signal of many inactivated ChrX genes reassures that the biallelic signal monitored in ChrX is unlikely to reflect a significant contamination from genomic DNA or cell-free RNA during library preparations. Additional file 3: Table S2 lists the supportive iSNPs for all the analyzed chromosomes in fibroblasts including the imprinted gene set.

### Identifying escapees in single cell primary fibroblasts

In the dataset of the primary fibroblasts, there are 232 and 485 genes that are supported by iSNPs evidence for ChrX and Chr17, respectively. As these cells lack information of genome phasing (Fig. 1b), information on escaping from X-chromosome is limited to the set of biallelic iSNPs (see Fig. 1b). We aggregated the iSNPs according to their corresponding genes (Fig. 1b). The aggregation is performed across different single cells and across multiple iSNPs within a specific cell-gene pair. A gene will be labeled escapee candidate when it is associated with multiple biallelic iSNPs. Altogether we identified 24 such genes (Table 1) which account for 10.3% of all expressed genes in ChrX. As expected, the fraction of genes on Chr17 showing biallelic expression is substantially higher (49.3%, Additional file 4: Table S3).

**Table 1.** Escapees from 101 primary single cell fibroblasts. A full list of all genes is available in Additional file 3: Table S4.

| Gene symbol | Gene type | # hSNPs per gene | Total iSNPs | # Cells with any observation | # Cells with biallelic observation<sup>a</sup> | Biallelic iSNPs Ratio<sup>b</sup> | Evidence poor<sup>c</sup> |
| --- | --- | --- | --- | --- | --- | --- | --- |
| CD99 | PAR | 28 | 248 | 101 | 97 | 0.435 |  |
| LAMP2 |  | 8 | 186 | 100 | 2 | 0.011 |  |
| RBM3 |  | 7 | 110 | 100 | 3 | 0.027 |  |
| ZFX |  | 36 | 456 | 100 | 80 | 0.226 |  |
| MTRNR2L10 |  | 10 | 699 | 98 | 80 | 0.200 |  |
| LOC550643 | ncRNA | 5 | 99 | 94 | 2 | 0.020 |  |
| SLC25A6 | PAR | 1 | 88 | 88 | 50 | 0.580 |  |
| C1GALT1C1 |  | 5 | 246 | 85 | 2 | 0.012 |  |
| SMC1A |  | 2 | 66 | 64 | 18 | 0.288 |  |
| DHRX | PAR | 60 | 134 | 63 | 12 | 0.097 |  |
| FAM104B |  | 17 | 937 | 63 | 2 | 0.002 |  |
| TSPAN6 |  | 1 | 61 | 61 | 4 | 0.066 |  |
| MAP7D3 |  | 9 | 127 | 59 | 3 | 0.024 |  |
| XIAP |  | 24 | 97 | 56 | 1 | 0.031 | * |
| HDHD1 |  | 19 | 186 | 49 | 8 | 0.091 |  |
| DDX3X |  | 9 | 99 | 41 | 6 | 0.091 |  |
| CA5BP1 | ncRNA | 18 | 77 | 39 | 2 | 0.039 |  |
| ZRSR2 |  | 11 | 42 | 34 | 6 | 0.143 |  |
| RBM41 |  | 16 | 63 | 29 | 5 | 0.079 |  |
| MST4 |  | 4 | 41 | 21 | 6 | 0.171 |  |
| JPX | ncRNA | 1 | 14 | 14 | 5 | 0.357 |  |
| KLHL4 |  | 15 | 27 | 9 | 1 | 0.074 | * |
| USP27X |  | 1 | 4 | 4 | 2 | 0.500 | * |
| RAB41 |  | 2 | 2 | 1 | 1 | 1.000 | * |

<sup>a</sup>The number of cells supporting the reported observations from all cells that have any observation; <sup>b</sup>Fraction of biallelically expressed iSNPs among all iSNPs associated with the gene, for example in USP27X there were overall 4 iSNPs, two of them were biallelic so the iSNPs ratio is 0.5; <sup>c</sup>Evidence poor applies for <10 iSNPs, or evidence based on one cell.

Table 1 lists the escapee candidates at varying degrees of support. For example, *ZFX* (Zinc finger X-chromosomal protein) and *SMC1A* (Structural maintenance of chromosomes protein 1A) genes are strongly supported with 103 and 19 biallelic iSNPs, respectively. A further increase in the reliability of identifying escapees is based on having at least 2 independent cells that contributed information on biallelic expression. We show that 21 out of 24 genes met this strict criterion (Table 1). Notably, among the identified escapees we detected only three PAR genes (*SLC25A6, CD99*, and *DHRSX*, Table 1). The assignment of these genes as escapees agrees with the expected PAR expression. From the number of biallelic PAR genes out of the expressed PAR genes, we estimated the false negative discovery rate for escapees to be as high as 70% (i.e., missed 7 of 10 expressed PAR genes). Additional file 4: Table S3 shows the support for Table 1.

### Quantifying allelic expression from clonal phased lymphoblasts

A major limitation in the protocol described above concerns the lack of parental haplotype phasing. Under this setting, iSNPs cannot be assigned to Xa or Xi. Consequently, the random choice of Xi which characterizes primary cells limits the discovery rate for escapees. We expanded the analysis of scRNA-Seq to female-originated lymphoblasts from the clonal cell-line GM12878 [38]. For this cell line, the sequence of the diploid genome is available with a reference for the paternal and maternal haplotypes [39]. Using the diploid genome as a reference, we definitively assigned the expression of sequenced reads covering hSNPs to either Xi or Xa. The availability of both parental haplotypes eliminates the biases that are associated with a single reference genome [40, 41]. In addition to the benefit of having two reference haplotypes, data from clonal cells drastically reduce the cell to cell variability that dominates primary cells. See also Additional file 1: Fig S4.

Figure 3a shows the expression profile for clonal lymphoblast single cells (n=25) (Additional file 2: Table S1, Additional file 5: Table S4). In any single cell, the monoallelic expression reflects the combination of an under-sampling of transcripts and the phenomenon of transcriptional bursting [28, 32, 33, 35]. It is clear that the maternal expression from the Xa dominates (Fig. 3a, top). An observation that agrees with the reported maternal Xa origin of cell-line GM12878 [38]. In most of the analyzed cells, a small but substantial fraction of the measured total expression is from the paternal, Xi chromosome (Fig. 3a, top). In contrast, Chr17 and the autosomal chromosomes show an equal expression from both alleles (Fig. 3a, middle and bottom panels).

**Fig. 3.**
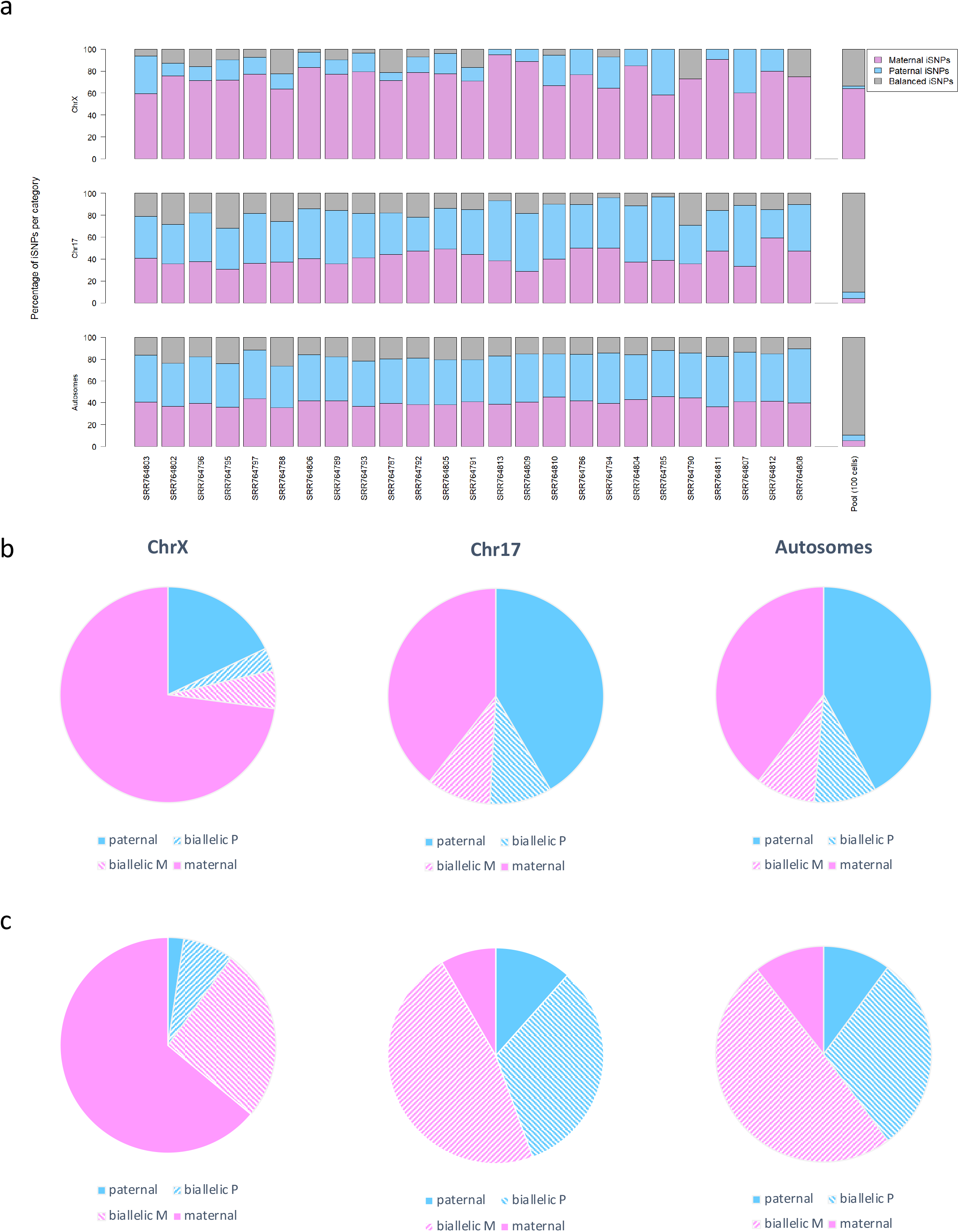
Quantifying the iSNPs’ labels from 25 single cell lymphoblasts. **(a)** Each single cell is partitioned according to its tagged allelic iSNPs on ChrX, Chr17, and all autosomal chromosomes. The iSNPs are associated with maternal (pink), paternal (light blue) and balanced expression (gray). The cells are ordered from left to right according to their iSNP contributions (Additional file 1: Fig. S4). On the right, the summary statistics of the Pool100 is shown. **(b)** A summary of the partition of iSNPs tags for all 25 single cells on ChrX, Chr17 and autosomal chromosomes. **(c)** A summary of the partition of iSNPs tags for Pool100. Blue and pink colors are associated with the paternal and maternal alleles, respectively. The striped pattern indicates biallelic iSNPs leaning towards paternal (blue) or maternal (pink) alleles. For single cells, the data is based on 375 iSNPs for ChrX, 808 iSNPs for Chr17 and 20,212 iSNPs for autosomal chromosomes. The data of Pool100 is based on 211 iSNPs for ChrX, 216 for Chr17 and 5,360 iSNPs for autosomal chromosomes. For the source data see Additional file 5: Table S4.

From the figure, it is evident that the phenomenon of transcriptional burst affects all chromosomes including ChrX. For assessing the impact of this phenomenon on identifying genes as escapees, we compared single cells with respect a pool of cells (Pool100, Fig. 3a, right bar). While the majority of the iSNPs from Chr17 display biallelic profiles, ChrX remains dominated by a maternal monoallelic expression.

Figure 3b is an aggregated view of ChrX, Chr17 and the autosomal chromosomes. The data are based on 375, 808 and 20,212 expressed hSNPs, respectively. Figure 3b (middle and right panels) shows an equal partition of the parental alleles from Chr17 and autosomal chromosomes (Fig. 3b, top). Performing the same analysis on data collected from Pool100 (Fig. 3c) shows that the partition of the parental alleles remains practically unchanged (compare the fraction occupied by pink and blue colors, Figs 3b-3c). Additionally, we observed a shift from a monoallelic (Figs 3b-3c, filled color) to a biallelic expression (Figs 3b-3c, stripped color). The fraction of the biallelic expression for Ch17 increased from 19% in single cells to 80% in Pool100, and for autosomal chromosomes from 18% to 79% (Fig. 3c, middle and right panels). The results from Pool100 indicate that the monoallelic expression observed in single cells is practically abolished by averaging the signal.

The results from ChrX (Fig. 3b (left) are fundamentally different relative to Chr17 or the autosomal chromosomes (Figs 3b-3c). The most notable difference is that only 21% of the expressed iSNPs are associated with the paternal Xi allele in ChrX (Fig. 3b, top). Furthermore, in analyzing Pool100, the fraction of biallelic expression remains bounded (a shift from 9% in single cells to 34% in Pool100). The observed pattern of ChrX from Pool100 (Fig. 3c, left) is best explained by an averaging of the stochastic monoallelic signal (at the same degree as the other chromosomes) while maintaining a strong signal of the Xa monoallelic expression. The strong evidence of X-chromosome inactivation in the single cells reassures that the observations of biallelic signals in ChrX are unlikely to reflect genomic DNA or cell-free RNA contaminations. See Additional file 5: Table S4 for lymphoblasts allelic ratio of all tested chromosomes and the Pool100.

### Identifying escapees from single cell lymphoblasts

Figure 4a is a gene-centric view that shows the iSNP allelic partition from lymphoblasts (colored according to their origin as maternal, paternal or mixed expression, see Methods). Only the subset of genes that are supported by multiple iSNPs is listed according to their ordered along the chromosomes. Altogether we report on 93 annotated genes on ChrX (Fig. 4a, 30 escapees and 63 inactivated genes). Note that the X-inactivated genes account for genes which are expressed primarily from the maternal Xa. A cluster of genes with a paternal expression at the tip of ChrX p-arm represents the expected biallelic expression from the PAR genes (Fig. 4a). Additional evidence for paternal expression is localized to the XIC with genes such as *XIST, JPX*, and *FTX*. While most of the escapees are supported by a limited number of iSNPs, a few of them such as *ZFX, CD99*, and *SLC25A6* are supported by a relatively large number of supporting iSNPs (48, 38 and 34, respectively).

**Fig. 4.**
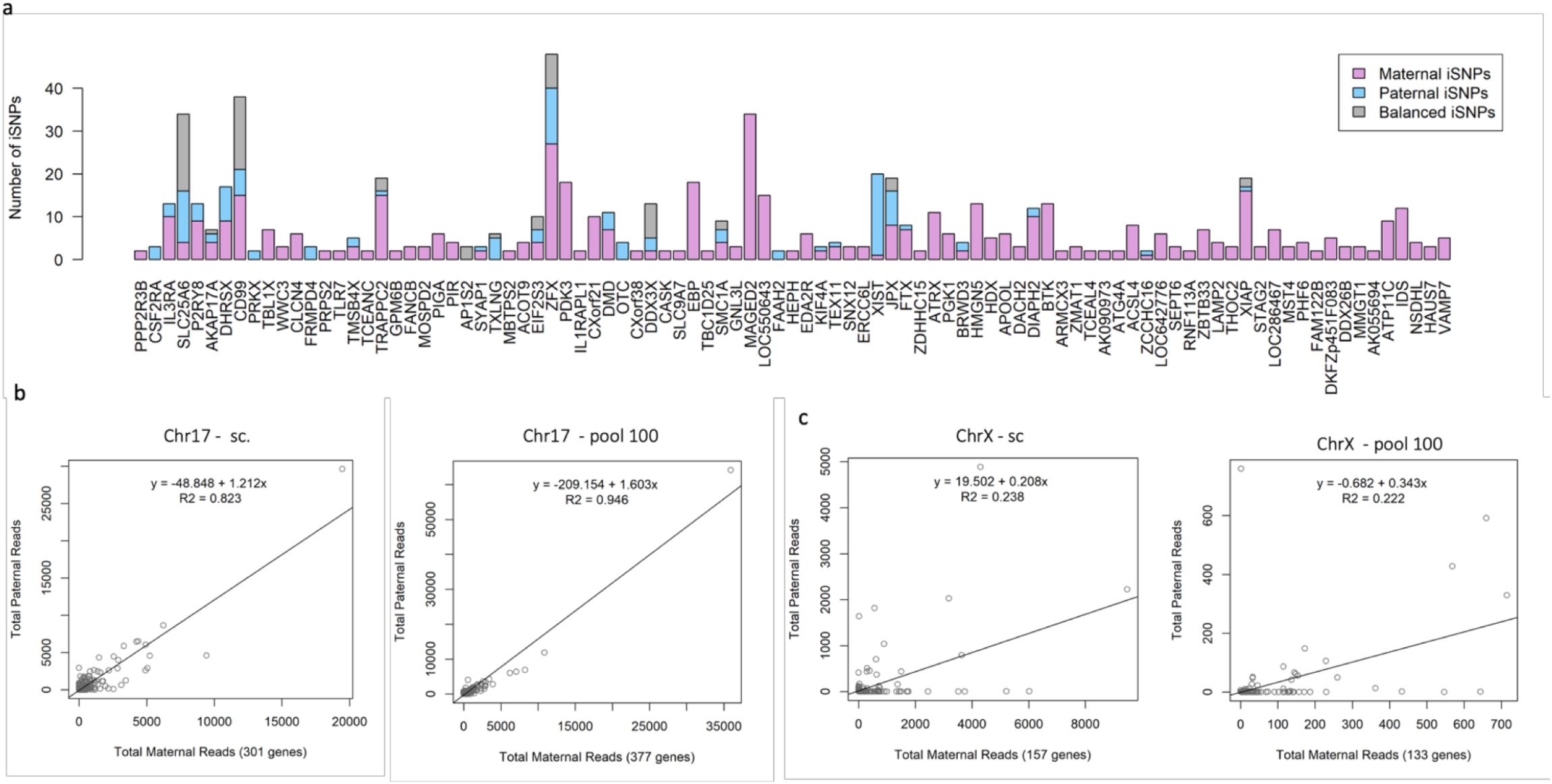
A gene-centric partition of alleles from lymphoblast cells. **(a)** For each gene on ChrX, the iSNPs parental partition is shown along with the number of iSNPs. For clarity, only genes that are supported by >=2 iSNPS are listed. A total of 93 genes in ChrX are listed by their order on the chromosome. The color code is according to the iSNP labels as paternal, maternal and balanced expression. For the source data, see Additional file 5: Table S4. **(b-c)** Correlation between the expression levels from the paternal and maternal alleles. The scatter plots show the expression levels of genes by the number of reads associated with maternal (x-axis) and paternal (y-axis) alleles. The number of analyzed genes for each scatter plot is indicated (on the x-axis, in parenthesis). Data shown are from Chr17 **(b)** and ChrX **(c)** based on single cells and Pool100. Note that the number of reads for the Pool100 data is 10-fold less with respect to the cumulative data extracted from single cells. For the source data, see Additional file 4: Table S3.

An alternative method for assessing the extent of the phenomenon of X-inactivation is by quantifying the evidence directly from the sum of all sequenced reads (abbreviated as the read-based protocol). Figures 4b-4c compare read counts from Chr17 (Figs 4b) and ChrX (Figs. 4c) by the paternal versus maternal origin. We compare the expression data from the single cells and the Pool100. A linear regression for the gene expression from Chr17 shows a high correlation fit-line (r^2^=0.823, Fig. 4b). As expected, the correlation is stronger in the data originated from the Pool100 (r^2^=0.946, Fig. 4b). We concluded that despite the monoallelic expression due to the transcriptional bursting phenomenon, a balanced allelic expression of all genes is strongly supported. For ChrX however, the resulting linear regression of the single cells is poor (r^2^=0.238, Fig. 4c), and was not improved by the data from the Pool100 (r^2^=0.222, Fig. 4d). Inspecting the expression data for ChrX shows that the regression lines actually lean toward the maternal Xa expression (x-axis). The expression data are consistent with two distinct regression lines for ChrX. One that matches the inactivated genes (parallel to the x-axis), and the other matches a trustfully biallelic expression. We conclude that the data representation (Fig. 4) accurately recaptured a gene-centric view of the X-chromosome inactivation phenomenon.

Applying the conservative iSNP-based protocol leads to the identification of 30 genes as escapee candidates that are also supported by the read-based protocol (Table 2). The read-based (i.e., labeling a gene as escapee by having a minimal number of paternal reads, see Methods) protocol expanded the escapee candidate list to include overall 49 genes (Additional file 4: Table S3).

**Table 2.** Escapees from 25 clonal single cell lymphoblasts.

| Gene symbol<sup>a</sup> | Gene Type | Total iSNPs | iSNPs Ratio<sup>b</sup> | # cells with any observation | # cells with paternal observation | Total Reads | Paternal reads | Paternal Ratio<sup>c</sup> | Evidence poor<sup>d</sup> |
| --- | --- | --- | --- | --- | --- | --- | --- | --- | --- |
| SLC25A6 | PAR | 34 | 0.882 | 18 | 16 | 9182 | 4887 | 0.532 |  |
| ZFX |  | 48 | 0.438 | 21 | 12 | 11697 | 2221 | 0.190 |  |
| CD99 | PAR | 38 | 0.605 | 17 | 10 | 5205 | 2027 | 0.389 |  |
| DDX3X |  | 13 | 0.846 | 12 | 10 | 1003 | 367 | 0.366 |  |
| XIST | ncRNA | 20 | 0.950 | 10 | 10 | 1652 | 1640 | 0.993 |  |
| JPX | ncRNA | 19 | 0.579 | 10 | 8 | 1931 | 1039 | 0.538 |  |
| EIF2S3 |  | 10 | 0.600 | 10 | 6 | 1322 | 706 | 0.534 |  |
| DHRX | PAR | 17 | 0.471 | 8 | 5 | 2379 | 1815 | 0.763 |  |
| SMC1A |  | 9 | 0.556 | 9 | 5 | 807 | 511 | 0.633 |  |
| TRAPPC2 |  | 19 | 0.211 | 12 | 4 | 4427 | 790 | 0.178 |  |
| DMD |  | 11 | 0.364 | 6 | 4 | 1533 | 160 | 0.104 |  |
| AKAP17A | PAR | 7 | 0.429 | 7 | 3 | 722 | 438 | 0.607 |  |
| AP1S2 |  | 3 | 1.000 | 3 | 3 | 99 | 54 | 0.545 |  |
| XIAP |  | 19 | 0.158 | 6 | 3 | 1944 | 437 | 0.225 |  |
| P2RY8 | PAR | 13 | 0.308 | 5 | 2 | 843 | 454 | 0.539 |  |
| TMSB4X |  | 5 | 0.400 | 5 | 2 | 372 | 70 | 0.188 |  |
| FAAH2 |  | 2 | 1.000 | 2 | 2 | 35 | 35 | 1.000 |  |
| DIAPH2 |  | 12 | 0.167 | 7 | 2 | 835 | 83 | 0.099 |  |
| CSF2RA | PAR | 3 | 1.000 | 1 | 1 | 58 | 56 | 0.966 |  |
| IL3RA | PAR | 13 | 0.231 | 6 | 1 | 837 | 120 | 0.143 |  |
| PRKX |  | 2 | 1.000 | 1 | 1 | 36 | 36 | 1.000 | * |
| FRMPD4 |  | 3 | 1.000 | 1 | 1 | 119 | 118 | 0.992 |  |
| SYAP1 |  | 3 | 0.333 | 2 | 1 | 190 | 57 | 0.300 |  |
| TXLNG |  | 6 | 1.000 | 1 | 1 | 197 | 170 | 0.863 |  |
| OTC |  | 4 | 1.000 | 1 | 1 | 44 | 44 | 1.000 | * |
| KIF4A |  | 3 | 0.333 | 3 | 1 | 265 | 27 | 0.102 | * |
| TEX11 |  | 4 | 0.250 | 3 | 1 | 244 | 84 | 0.344 | * |
| FTX | ncRNA | 8 | 0.125 | 3 | 1 | 716 | 35 | 0.049 | * |
| BRWD3 |  | 4 | 0.500 | 3 | 1 | 88 | 48 | 0.545 | * |
| ZCCHC16 |  | 2 | 0.500 | 1 | 1 | 57 | 32 | 0.561 | * |

<sup>a</sup>Only 30 genes that are supported by iSNP-based and read-based methods are listed. A full list of all genes (49) is available in Additional file 3: Table S4. <sup>b</sup>Fraction of iSNP supporting the paternal or biallelic observations out of total recorded iSNPs for a gene, for example in ZCCHC16 there were overall 2 iSNPs, one of them was paternal so the iSNPs ratio is 0.5. <sup>c</sup>Fraction of parental reads out of total read for a gene. <sup>d</sup>Evidence poor apply for genes with a cell-based inconsistency or having <50 paternal reads with only one supporting cell.

Testing the parental origin of alleles along a gene in the same cell is a strict test for the reliability of the iSNPs. This test is only valid for genes with multiple hSNPs. Such genes that are supported with two or more expressing hSNPs account for 44% of the genes. We consider a gene to be consistent if the expression along the gene in a specific cell is not monoallelic to both alleles. Altogether, we identified 3 inconsistent genes - *TEX11, FTX*, and *ZCCHC16*. For another 6 genes, the inconsistency is only partial as there are other observations of biallelic expression. The estimate from full inconsistency (3 out of the 29 genes that were eligible for this test) suggests that an upper limit for a faulty interpretation of 10%. Additionally, the iSNP-based protocol identified 9 out the 11 expressed PAR genes. Thus, we extrapolate the escapee detection rate to be 82%. Interestingly, analyzing Chr17, under the assumption that there is no systematic allelic bias [36], showed that 7.3% and 9.6% of the genes were associated with maternal and paternal monoallelic expression, respectively. These results provide an upper limit of 17.9% to the likelihood of false gene labeling in Chr17 and can be used for estimating the limitation of the method.

Demanding biallelic evidence from at least two cells reduces the number of escapees from 49 to 18 (including 5 PAR genes). Table 2 shows the discovered escapee candidates along with their supporting evidence (Table 2).

We anticipate that a larger number of cells with scRNA-Seq data, deeper sequencing with higher coverage, and samples that present a denser map of hSNPs (e.g., from remote ethnic groups) will substantiate the annotation confidence of escapee genes (Table 2).

### Comparison of the identified escapees to current knowledge

Escapees are estimated to occupy 15-25% of all ChrX genes in humans [13]. A unified catalog was compiled by [13] from the integration of four independent studies that covers 1144 genes from ChrX. The genes in this catalog are manually partitioned into nine defined categories (see Methods). The largest one accounts for genes that lack information (45%) [13]. About 15% of the genes (168/1144) are considered ‘escapee-associated’ (See Methods). Among them, only 51 genes (4.5%) show a consistent expression from Xi in various experimental settings and across different methods.

We tested the correspondence between the identified escapees from our study and this literature-based catalog [13, 24]. To this end, we applied a hypergeometric statistical test (see Methods) to assess the overlap of the different escapee gene lists (Fig. 5). We consider the compiled set of ‘escapee-associated’ genes as a gold standard to test escapee’s discovery rate in our study (total of 124 genes, excluded PAR genes, collectively called Balaton-Esc).

**Fig. 5.**
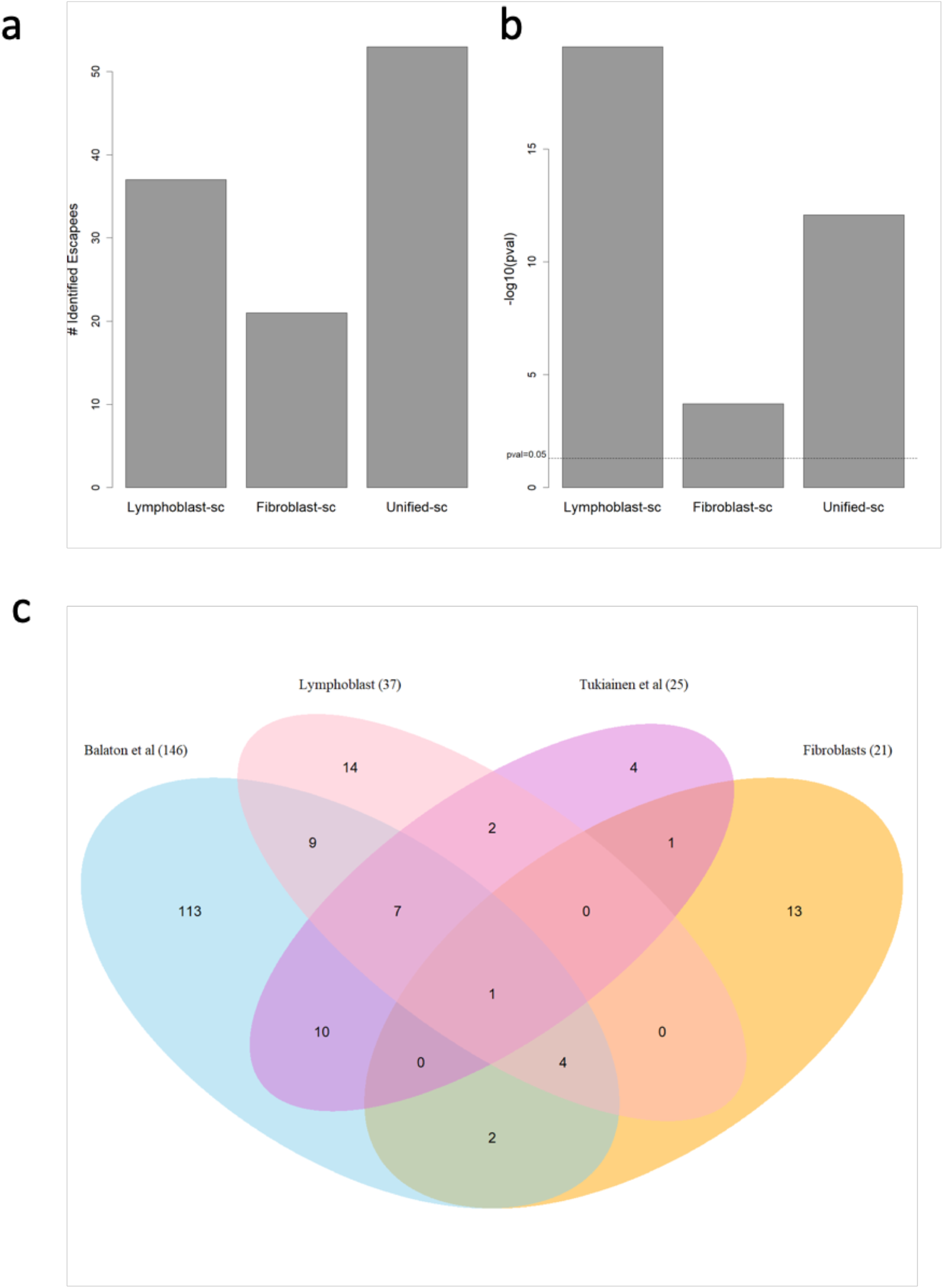
Identified escapees and statistical significance of the overlap with literature-based catalog compiled by Balaton et al. [13]. **(a)** The numbers of escapees identified by each of the analyses. The numbers include only genes that were present in Balaton et al. [13] and exclude PAR genes. **(b)** Statistical analysis based on the hypergeometric distribution measuring the overlap between the literature-based list as presented by Balaton et al. [13] and the escapee assigned in this study (as in (a)). Y-axis is the -log10(x) of the calculated p-value. **(c)** Venn diagram of the 4 sets of escapees according to the analyzed fibroblasts and lymphoblasts, the Balaton-Esc collection [13], and the Tukiainen-Esc [16]. Including PAR. For details see text. Source data in Additional file 6: Table S5.

Figure 5a shows the number of identified escapees from the fibroblasts and lymphoblasts (excluding PAR genes). Note that only genes that are included in the Balaton-Esc benchmark are included in this analysis (Fig. 5). Figure 5b shows the statistical significance of the overlap between the gene lists from Fig. 5a and the Balaton-Esc [13]. As can be seen, there is a significant overlap between the escapees from lymphoblasts (Table 2 and Additional file 4: Table S3) and Balaton Esc list (Fig. 5b, p-value = 7.43E-8). Applying the same test for the primary fibroblasts (Table 1) resulted in a lower significance (p-value = 4.07E-2).

Figure 5c depicts the overlap genes between the escapees identified in our study and the Balaton-Esc catalog (168 genes including PAR genes) [24]. We also included a complementary resource based on 940 transcriptomes from scRNA-Seq (25 escapee genes, Tukiainen-Esc) [16]. The Venn diagram shows that each of the above studies contributes to the current knowledge on escapees. Escapees from the two external resources overlap by 18 out of 25 reported genes (72%). As shown in Fig. 5c, 62% of the escapees reported from the lymphoblasts overlap with the external escapee lists, while the fibroblasts are supported by only 38% overlap. Notably, most of our discovered candidate escapee genes from fibroblasts (62%) have no correspondence with the other tested lists (Additional file S6: Table S5).

### LncRNAs extend the list of escapee candidates

Most studies on escapees focus on the coding genes. However, our protocol for escapee discovery is based on whole-genome alignment, and thus it is suitable also for other ncRNAs (Additional file 7: Table S6). As ChrX has a high density of short and long ncRNAs [3] (Fig. 1a), we search for long non-coding RNAs (lncRNAs) among the expressed hSNPs that meet our escapee criteria (Fig. 6). To the best of our knowledge, it is the first systematic and unbiased effort for mapping lncRNA escapees. Altogether we identified 15 lncRNAs as escapee candidates, among them only a few were previously studied. The location of the lncRNAs and coding escapes along ChrX is shown (Fig. 6a). We tested the positions of escapees along the ChrX relative to all ChrX genes. While the positional distribution for lncRNA escapees is similar (Kolmogorov–Smirnov test, p-value = 0.57). it is different for coding escapees (Kolmogorov–Smirnov test, p-value = 0.004). This is in accord with the overdensity of coding escapees at ChrX p-arm [17, 21, 22] (Fig. 6a).

**Fig. 6.**
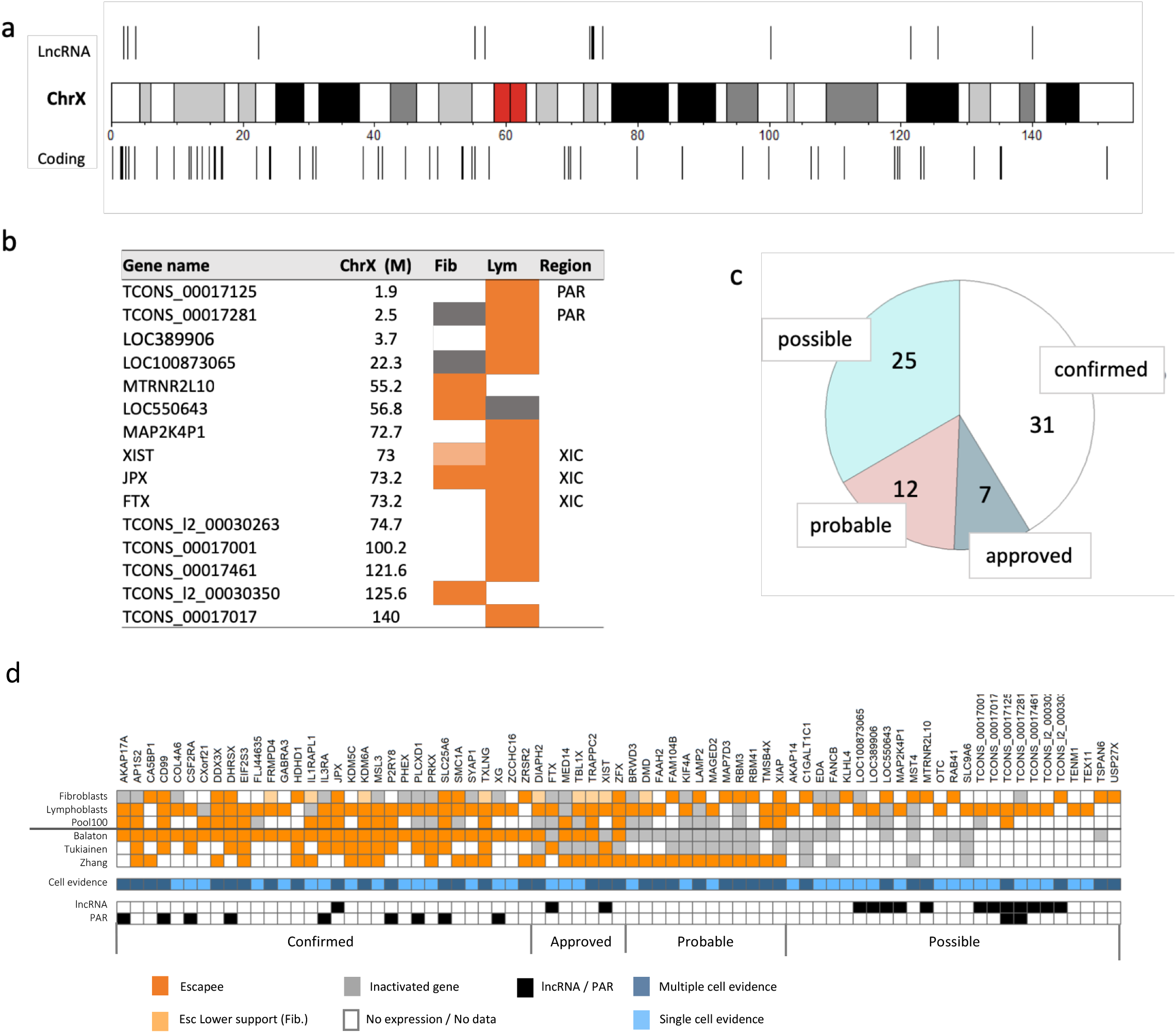
LncRNAs assigned as escapees, and the groups of escapees according to their confidence level. **(a)** The dispersal of escapees along ChrX. Escapees belong to the lncRNAs and to the coding genes are indicated above and below the schematics of ChrX, respectively. **(b)** A table listing the 15 lncRNA escapee candidates. The orange color indicated escapee and the gray inactivated. An assignment that is based on a single iSNP is labeled with light orange. The white indicates lack of report or no expression. XIC, X inactivation center region. **(c)** Partition of the 75 genes that were mentioned as escapee candidates in this study. The categories are labeled ‘confirmed’, ‘approved’, ‘probable’ and ‘possible (see text). The two external resources that are used to define the groups are from the literature [13] and from single-cell by [16]. For a group of ‘probable’ escapee, we used escapee’s annotation from [10] as evidence. Note that 11 of the 15 listed lncRNA genes are included in the ‘possible’ escapee set. **(d)** A summary of the evidence-based groups for 75 genes. Genes are sorted according to the 4 evidence groups (as in **c**) The escapees are colored orange. Light orange indicates escapees that are supported by a single evidence from one cell only in fibroblasts. Inactivated genes are colored gray. White color indicates no expression or lack of report. The cell evidence is color coded showing support by one (light blue) or multiple cells (dark blue). Cases where evidence are based on Pool100 only are also marked light blue. PAR genes and lncRNAs are marked. Source data is in Additional file 6: Table S5.

Figure 6b lists all 15 identified lncRNA escapee genes, among them, are ncRNA genes from the XIC that coordinate the activation and maintenance of X-inactivation. Many of the lncRNAs are localized at transcriptionally active segments (e.g., within the PAR or the XIC), while others are localized in non-conserved regions which are enriched with long and short ncRNAs. For additional lncRNAs, including inactivated genes see Additional file 7: Table S6.

### Evidence-based partition of escapee genes

Figure 6c summarizes the partition according to the evidence for all genes that are reported with any levels of confidence as escapees (Additional file 6: Table S5). This list includes 75 candidates that are reported in this study, including evidence from Pool100 and a collection of novel lncRNAs. Fig. 6d provides a detailed list of the finding from this data in view of serval external resources. The number of cells providing evidence is also indicated for every gene.

The comparison to the cumulative external resources enables us to roughly estimate the upper bound for the false positives rate of both datasets (Additional file S6: Table S5). The number of escapees that agrees between the lists of each cell type and the labels according to the Balaton catalog [13] was calculated. Solely based on these comparisons, we estimate that the fibroblasts false positive rate is bounded to 52%, while for the lymphoblasts, it is bounded to 30%. This is, of course, an upper bound since it is possible that not all escapee genes are included in the Balaton catalog [13]. Additionally, most of the studies compiled in Balaton-Esc catalog have used lymphoblasts and not fibroblasts (See Methods).

Taking external resources into account, we matched each gene according to the quality of the independent support associated with it (Fig. 6c and 6d). Specifically, we partitioned the 75 gene list to four groups: (i) Genes are labeled “confirmed” if they are reported as escapees by both previously discussed external resources [13, 16]. There are 31 such genes, for which this study provides a further confirmation for their identity as escapees. (ii) Additional 7 genes are labeled “approved”. These genes are tagged as escapees by only one of the two external resources [13, 16]. For these genes, the independent evidence from this study approves their identity. (iii) Additional 12 genes are marked as “probable” escapees. These genes are assigned according to the agreement with an additional external report reporting on 114 escapees [10], which was not included in the literature-based catalog [13], and thus can be considered as an independent resource. (iv) Additional 25 genes are marked as “possible” escapees. These genes lack any literature evidence for supporting their identity, thus their assignment as escapees remains less supported. In this set, there are 15 genes that were not reported by any of the three discussed external resources [10, 13, 16]. The majority of the overlooked lncRNAs belong to this group. These genes are awaiting additional independent evidence for established their assignment. The source data for Fig. 6d is in Additional file 6: Table S5.

### Revising annotations of escapees

The largest group from our identified set fully agrees on the annotation provided with the two external resources (‘confirmed escapees’, Fig. 6c). While we will not discuss this most validated gene set, it is important to mention that among the 31 ‘confirm’ escapees, 9 were repeatedly validated as PAR genes and other are established escapees (Fig. 6d). Two overlooked escapee lncRNAs (TCONS_00017125, and TCONS_00017281, Fig. 6b) are located at the PAR region that is exceptionally active in biallelic transcription. Due to the lack of any evidence from external resources they were assigned among the set of ‘possible’ escapees. We anticipate that these lncRNAs belong to the other biallelically expressed PAR genes.

The gene groups that are marked as ‘approved’ (7 genes) and ‘probable’ escapees (12 genes, Figs, 7c-7d) are reported by some external resources but not others, indicating inconsistency in their assignment. Our findings help to resolve some of these conflicts (total 19 genes).

Two of the genes (*XIST* and *FTX*) belong to XIC and are critical to the X-inactivation process [3, 42] and have conflicting annotations. Actually, we observed that the strongest expressed gene from the Xi in lymphoblasts is *XIST* (Additional file 4: Table S3). We also identified *XIST* with some evidence of biallelic expression among primary fibroblasts. We argue that it was mistakenly labeled by [13] as an inactivated gene. We also have monitored *FTX* expression from Xi from both our datasets, in agreement with evidence from [16]. This might indicate that the escapee signal of lncRNAs is more easily detected at a single-cell resolution. We proposed to revise the annotations for these genes.

The two external resources [13, 16] indicated *XIAP* as inactivated. In contrast, we have identified it in both fibroblasts and lymphoblasts as a true escapee (Fig. 7d), which was further supported in [10]. Interestingly, inconsistency in the annotation of *XIAP* was noted in individuals from different ethnic groups [10]. The following ‘approved’ escapees (*TBL1X, TRAPPC2, ZFX, DIAPH2*) were listed in the Balaton-Esc and were also supported by [10]. Based on our supportive evidence, such partial inconsistency is resolved.

The 12 ‘probable’ escapees corroborate findings from [10] in which lymphoblasts from different ethnic origin were tested by analyzing their expressed hSNPs. Our results support the findings for all 12 ‘probable’ escapees (*TMSB4X, DMD, RBM3, MAGED2, FAM104B, FAAH2, KIF4A, BRWD3, RBM41, LAMP2, XIAP*, and *MAP7D3*). While most of these genes have a substantial support *BRWD3, PRKX* and *XIAP* are marked as ‘evidence poor’ (Tables 1-2). Nevertheless, a comparison to a study of Klinefelter syndrome (47, XXY) individuals from the Danish population reveals some interesting compatibilities. The individuals in the study were tested for sex biases from ChrX. It was found that 16 genes were overexpressed. Among them, *XIST*, numerous PAR genes as well as a few others were confirmed in this study as escapees (*EIF2S3, PRKX*) [43]. Specifically, *TMSB4X* (a ‘probable’ escapee) was associated with ChrX dosage effect in Klinefelter syndrome, presumably due to its escapee characteristics.

### A small set of constitutive escapees

Only 4.5% of all known ChrX genes are unquestionable escapees that are consistently detected by complementary methods and are considered ‘constitutive escapees’ according to [13]. Within our two cell types analyzed datasets there are 5 escapee genes (*DDX3X, ZFX, SMC1A, JPX, and XIAP*) in common which confirm their strong tendency to escape X-inactivation (Tables 1-2, and Additional file 3: Table S4, Additional file 6: Table S5). Four of these genes (ZFX, *JPX, SMC1A*, and *DDX3X*) were also reported by others as constitutive escapees. These constitutive escapees act within the nuclei in transcriptional regulation (*DDX3X* and *ZFX*) and in chromosomal dynamics (*SMC1A* and *JPX*). We propose that *XIAP* belongs to this small set of escapees showing a robust characteristic across different cells. In a study on a large population of Klinefelter syndrome individuals, *XIAP* was identified among the few ChrX overexpressed genes [43].

Having a robust and reliable list of escapees is critical in understanding sex-dependent phenotypic variability. In addition, the profile of escapees may explain the clinical severity of individuals with X-chromosome aneuploidy (e.g., Turner (XO) and Klinefelter (XXY) syndrome) [43, 44]. The expression level of certain escapee genes was also attributed to a number of sexually dimorphic diseases (e.g., cancer) [26], autoimmunity [45] and more. Therefore, our results refine the list of the relevant escapees and expand it.

## Conclusions

In this study, we show that X-inactivation and escape from it can be successfully studied from single cells, including primary, untransformed cells. The cumulative information from individual cells increases the potential of identifying escapees and inactivated genes. Moreover, despite the use of a limited number of cells, clonal cells with genomic phasing are valuable for detecting escapees at high confidence. Generalizing the ASE based method to uncharacterized genomic loci resulted in for the first time, in a complete report on lncRNAs escapees. We affirm that even with a modest number of analyzed cells, the cumulative knowledge on escapees was reproduced, extended and numerous conflicting findings were resolved.

## Methods

### Outlook

The pipeline for escapee identification using allele quantification was performed over two independent single cell datasets: (i) A collection of 104 cells of primary fibroblasts from a female newborn (UCF_1014). Additional file 2: Table S1 shows the quantitative information associated with the analyzed 104 primary fibroblasts; (ii) A collection of 25 single cells of clonal GM12878 lymphoblast from female (NA12878, Additional file 2: Table S1). For GM12878 cells, the pipeline was also applied to a pool of 100 cells (Pool100).

The analysis of both datasets was performed for ChrX, Chr17 and all autosomal chromosomes. For each of the datasets, we used scRNA-Seq or RNA-Seq for Pool100 (see Additional file 1: Text). From additional genomics data, extensive lists of all available heterozygous SNPs (hSNPs) for the two datasets were compiled. These lists are candidate sites for informative allelic expression. A detailed description of the pipeline protocols is available in Additional file 1: Text. Fig. S1 that provides a scheme of the workflow for identifying escapees from scRNA-Seq data.

### Datasets

A dataset for scRNA-Seq of fibroblast UCF1014 was downloaded from the European Genome-phenome Archive (https://www.ebi.ac.uk/ega/home) using accession number EGAD00001001083. The data was produced by Borel et al. [28]. DNA-seq of UCF1014 was also downloaded from EGAD00001001084. The quality of the NGS data from scRNA-Seq in term of sequencing errors, PCR biases, and mapping errors was thoroughly discussed [28].

A dataset for scRNA-Seq of GM12878 cell-line from female lymphoblast NA12878 along with Pool100 was downloaded from the Gene Expression Omnibus (GEO) database (www.ncbi.nlm.nih.gov/geo) using the accession number GSE44618 [38]. The NA12878 phased genome was downloaded from http://sv.gersteinlab.org/NA12878_diploid/NA12878_diploid_genome_2012_dec16.zip. The SNP locations were downloaded from: http://sv.gersteinlab.org/NA12878_diploid/CEUTrio.HiSeq.WGS.b37.bestPractices.phased.hg19.vcf.gz.

For additional information and detailed protocols, see Additional file 1: Text.

### RNA-Seq alignment

RNA-Seq from single cells (scRNA-Seq) and pooled cells were aligned to a relevant reference genome. Datasets of scRNA-Seq from fibroblasts were aligned against UCSC hg19 reference genome after removal of the Y chromosome. The lymphoid scRNA-Seq and RNA-seq are aligned against the Paternal and Maternal haplotypes of the phased diploid NA12878 genome at the hg19 format. Reads aligned to hSNPs were counted such that each of the reads is resigned to one of the two alleles. Each hSNP, expressed in every single cell with a minimal number of 7 aligned reads was considered an informative SNP (iSNP). For details see Additional file 1: Text. Quantitative values of the number of iSNPs for each cell of the 104 primary fibroblasts are shown in Additional file 1: Fig. S2 and Additional file 2: Table S1. Quantitative values of the number of iSNPs for each cell of 25 single cell clonal lymphoblasts are in Additional file 1: Fig. S4 and Additional file 2: Table S1.

### Quantification and Annotation of Alleles by Expression

For the primary fibroblasts without phased haplotype data, the Allelic Ratio (AR) of each iSNP is determined as follows:

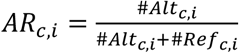

Where c indicates a cell and i indicates an iSNP. #Alt and #Ref refer to the number of reads aligned to the alternative and the reference allele, respectively.

For the lymphoblasts that are associated with a phased genome, the Allelic Ratio (AR) of each iSNP is determined as follows:

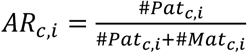

Where c indicates a cell and i indicates an iSNP. #Pat and #Mat refer to the number of reads aligned to the paternal and maternal allele, respectively.

An iSNP with AR of <=0.1 or AR >0.9 is considered monoallelic and an iSNP associated with 0.1<AR≤0.9 is considered biallelic.

For the fibroblasts data, tagging a specific iSNP as escapee relies on annotating multiple iSNPs of a gene as biallelic. For the clonal lymphoblasts data, the paternal ChrX is associated with Xi. Therefore, iSNPs with paternal or biallelic AR (expression from the paternal ChrX) are indicative of an escapee gene. An additional protocol for identifying escapees from the clonal single cell lymphoblasts is based on accumulating reads across cells with an escapee criterion of a minimal number of 7 paternal reads per gene. Lymphoblast Pool100 cells were quantified such that in each gene, the sum of hSNP reads with a paternal expression ratio of AR >0.1 indicates an escapee gene. For further details, the quantitative values of the resources, and the NGS tools that were used, see Additional file 1: Text (Sections 1-1.4).

### Cell outliers according to extreme biallelic expression

A possible cause of faulty interpretation for biallelic expression of a gene can originate from a single cell experiment that accidentally included two (or more) cells instead of one cell. Therefore, if for a specific cell, the proportion of biallelic expression shows an extreme value, the data associated from that cell could potentially be explained by having a mixture of two cells expressing two different Xi (coined a doublet). We have calculated for each cell the biallelic ratio as *biallelic ratio= biallelic iSNPs/total iSNPs*. Following this investigation, we have excluded three single cells which had a biallelic ratio of over 3 standard deviations from the average of the entire cells. After filtering these cells outliers, 101 single cells remained for further analyses.

As the lymphoblasts were handpicked. No such procedure was needed.

### A comparison to annotated ChrX gene catalog

There are 1144 known genes in ChrX that were annotated and compiled into a comprehensive catalog [13]. All genes were annotated and partitioned into 9 categories: (i) PAR, (ii) escapee, (iii) mostly-escapee, (iv) variable-escapee, (v) mostly-variable-escapee, (vi) discordant, (vii) inactivated genes, (viii) mostly-inactivated, and (ix) genes having no data. The set of criteria for assigning a gene to any of these categories was described in [13]. Annotation for the 9 categories is based on a careful analysis according to major publications combining numerous measurements for genes expression characteristics of ChrX [13]. The reports used for compiling the catalog were from four major resources extracted from [13, 17, 20, 46].

From the genes on ChrX, an “escapee associated” list was compiled from the 6 first categories (i-vi) associated with escapees, including PAR and discordant genes. This unified set corresponds to 15% of the genes, the rest are divided between inactivated-related categories (40%) and genes that are are annotated as “no data” (45%).

### Statistical analysis

Hypergeometric probability between our results and the external annotated catalog was calculated by comparing the correspondence of any two lists of escapees. We used standard notations of N, k, n and x: N symbolizes all genes on ChrX from [13] with a label other than “No data”; k is the number of escapees by [13] which are associated with any escapee annotations (i.e. the escapee-associated); n is the number of escapees we identified by any of the settings from our protocols; x the number of genes in our list that match the literature-based escapee list in k. P(x) is the probability that an n-trial will result in a value that is > x. See Additional file 1: Text (Section 1.5)

### Additional datasets for annotated escapees

Lists of the annotated escapees according to Tukiainen et al. [16] and Zhang et al [10] were downloaded from the publication supplemental materials. Accordingly, the status of escapee genes was analyzed across the external resources reported [10, 13, 16]. See Additional file 1: Text (Section 1.6)

## Abbreviations

Alt: Alternative allele
AR: Allelic Ratio
ASE: allele specific expression
Chr: chromosome
hSNP: heterozygous SNP
iSNP: informative SNP
lncRNA: long non-coding RNA
Mat: maternal
ncRNA: noncoding RNA
NGS: next generation sequencing
PAR: pseudoautosomal region
Pat: paternal
Ref: reference allele
RNA-Seq: NGS for RNA sequencing
Sc: single cell
SNP: single nucleotide polymorphism
Xa: X activated
Xi: X inactivated;
XIC-: X inactivation center.

## Acknowledgements

We thank Nati Linial for useful discussions and critical comments. We thank Liran Carmel, Shahar Shohat and Michal Linial’s lab members for a critical reading of the manuscript. We thank Yuval Nevo and the CSE system for technical assistance. KWK is a recipient of the Ariane de Rothschild Scholarship. This study used best practice protocols that are compiled in EU Elixir collection.

## Authors’ contribution

KWK and ML wrote the manuscript, designed and performed the experiments, and the data analyses. Both authors read and approved the final manuscript.

## Ethics approval and consent to participate

Not applicable.

## Availability of data

All data generated or analyzed during this study are included in this published article as supplementary material. The datasets are available according to the following sources:

Fibroblasts UCF_1014 DNA-seq from European Genome-phenome Archive respiratory dataset EGAD00001001083 (https://www.ebi.ac.uk/ega/datasets/EGAD00001001083)

Fibroblasts Single cells RNA-Seq from European Genome-phenome Archive respiratory dataset EGAD00001001084 (https://www.ebi.ac.uk/ega/datasets/EGAD00001001084)

Lymphoid genome of NA12878 from Gerstein Lab, Yale University, http://sv.gersteinlab.org/NA12878_diploid/NA12878_diploid_genome_2012_dec16.zip.

Lymphoid SNPs from Gerstein Lab, Yale University,

http://sv.gersteinlab.org/NA12878_diploid/CEUTrio.HiSeq.WGS.b37.bestPractices.phased.hg19.vcf.gz.

Lymphoid GM12878 single and pooled cells RNA-Seq from Gene Expression Omnibus, at http://www.ncbi.nlm.nih.gov/geo/query/acc.cgi?acc=GSE44618.

## Additional files

Additional file 1: 1.1-1.6 Detailed methods and protocols (Text)

Fig. S1. A workflow for identifying escapee genes from single cells RNA-Seq data

Fig. S2. Informative SNPs of 104 single cell primary fibroblasts

Fig. S3. The Biallelic ratio of 104 fibroblasts.

Fig. S4. Informative SNPs on 25 clonal female lymphoblasts

Additional file 2 Table S1: Quantitative data of primary fibroblast UCF_1014 cells and Identifiers and quantitative data of RNA-Seq for 25 lymphoid single cells and Pool100

Additional file 3: Table S2. A list of informative SNPs (iSNPs) along with their labeling on ChrX, Chr17, all autosomal chromosomes and imprinted genes from 101 fibroblast single cells.

Additional file 4: Table S3 Gene-centric tagging determined by iSNPs and a read-based method for the Fibroblast, Lymphoblast and Pool100 datasets on ChrX, Chr17 and all autosomal chromosomes.

Additional file 5: Table S4. A list of informative SNPs (iSNPs) along with their labeling on ChrX, Chr17 and autosomes from single cell lymphoblasts and Pool100 lymphoblasts.

Additional file 6: Table S5. Table comparing annotation of all 75 ChrX escape candidates.

Additional file 7: Table S6. LncRNAs informative SNPs (iSNPs) along with their labeling on ChrX on both fibroblasts, and Lymphoblasts.

