## Supplementary material for "Human Genes Escaping X-inactivation Revealed by Single Cell Expression Data": Sup file 1 (fig S1-S4)

**Additional file 1: Supplemental Materials and Methods (text)**

**Fig. S1.** A workflow for identifying escapee genes from single cells RNA-Seq data.
**Fig. S2.** Informative SNPs of 104 single cell primary fibroblasts
**Fig. S3. The** Biallelic ratio expression of fibroblasts.
**Fig. S4.** Informative SNPs on 25 clonal female lymphoblasts

### 1.1. Resources:

- Fibroblasts UCF_1014 DNA-seq: EGAD00001001083 (https://www.ebi.ac.uk/ega/datasets/EGAD00001001083)
- Fibroblasts Single cells RNA-seq: EGAD00001001084 (https://www.ebi.ac.uk/ega/datasets/EGAD00001001084)
- Lymphoid genome of NA12878 from Gerstein Lab, Yale University, http://sv.gersteinlab.org/NA12878_diploid/NA12878_diploid_genome_2012_dec16.zip.
- Lymphoid SNPs from Gerstein Lab, Yale University: http://sv.gersteinlab.org/NA12878_diploid/CEUTrio.HiSeq.WGS.b37.bestPractices.phased.hg19.vcf.gz.
- Lymphoid GM12878 single and pooled cells RNA-Seq from Gene Expression Omnibus http://www.ncbi.nlm.nih.gov/geo/query/acc.cgi?acc=GSE44618. Canonical transcripts list available from the UCSC known gene list [1].

### 1.2. Sequencing analytical procedures and tools:

#### 1.2.1. Alignment and genomic sequencing tools:

- BWA [2]
- STAR [3]
- SAMTools [4]
- Bowtie2 [5]
- LiftOver available at UCSC toolbox [6].

#### 1.2.2. SNP calling

- GATK best practices procedure [7].
- BCFtools utilities [8].

#### 1.2.3. Reads Cleaning

- Trimmnomatic [9].
- FASTX (http://hannonlab.cshl.edu/fastx_toolkit/index.html)

#### 1.2.4. Allele specific counting

- Allelcounter-master [10].
- HTSeq [11].

### 1.3. Statistical analysis and useful modules

- Rstudio
- R Core Team (2018). R: A language and environment for statistical computing. R Foundation for Statistical Computing, Vienna, Austria. URL https://www.R-project.org/.
- R ComplexHeatmap [12] package
- Biopython [13]

**A workflow for single cell genomics and transcriptomics analyses**

The following description applies to the workflow designed for two cell types.

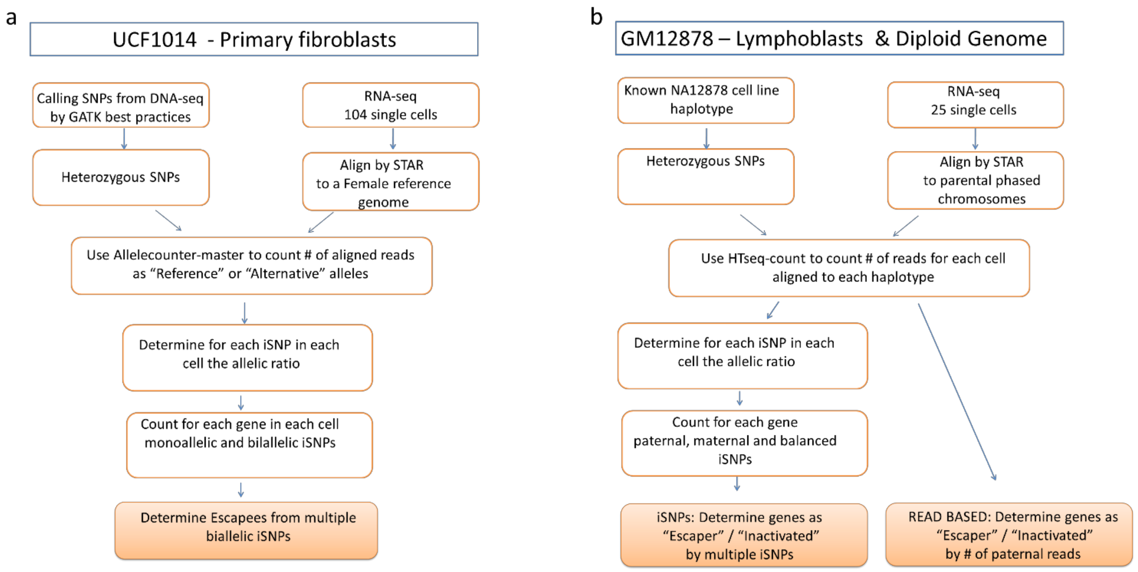


**Fig. S1.** A workflow for identifying escapee genes from single cells RNA-Seq data. **(a)** The protocol applied for identifying escapees using scRNA-seq from 104 UCF1014 primary fibroblast single cells coupled with their corresponding DNA-seq. Identical protocols were applied to ChrX, Chr 17 and the autosomes. **(b)** An outline scheme describing the protocols that were used for analyzing 25 single cell GM12878 lymphoblasts with phased haplotypes.

#### 1.3.1. Female primary fibroblasts

##### 1.3.1.1. Single cell primary fibroblasts and the reference genome

DNA-seq was based on female newborn primary fibroblast culture (called UCF_1014). Data were extracted from EGAS00001001009 study (https://www.ebi.ac.uk/ega/studies/EGAS00001001009) dataset EGAD00001001083 (https://www.ebi.ac.uk/ega/datasets/EGAD00001001083). For more details see [14, 15].

We realigned the 2 DNA-seq samples to the UCSC hg19 reference genome using BWA [2]. Variations were called using GATK best practice procedure [7]. In order to consider a SNP for further analysis, we request that the SNP appears in both VCFs (bcftools isec -n+2 -o UCF_1014.vcf -O v -p UCF_1014/ -w1) [8]. Only heterozygous loci in the VCF were compiled for further analysis (referred to as hSNPs).

##### 1.3.1.2. Single cell fibroblasts RNA-Seq

RNA-seq datasets of single cells were obtained from EGAS00001001009 study, dataset EGAD00001001084 (https://www.ebi.ac.uk/ega/datasets/EGAD00001001084) [15]. The preparation of the samples is described in [14].

For consistency, we chose to analyze only samples that were amplified under a similar amplification protocol (i.e., 22 cycles of PCR, total of 104 cells). The RNA-seq reads were preprocessed using Trimmnomatic [9] with the following parameters ILLUMINACLIP:NexteraPE-PE.fa:2:30:10 LEADING:3 TRAILING:3 SLIDINGWINDOW:4:15 MINLEN:50. Fragments from the scRNA-seq data were realigned to UCSC hg19 reference without the Y chromosome using STAR [3]with the default parameters. Repeated reads were marked using Picard, and RNA-seq was indexed using SAMtools [4]. The number of raw and aligned reads are shown in Additional file 2: Table S1.

##### 1.3.1.3. Single cell fibroblasts allelic imbalance analysis

All the reads from each BAM alignment file were assigned to the Reference or Alternative allele according to the SNPs on the VCF by Allelcounter-master [10]. For an informative SNP (iSNP) to be considered, an expression report of >=7 reads at the SNP location, in a single cell, was required. Replacing the predetermined read threshold by a cell-specific threshold that was calculated out of the aligned sum of reads for each cell, had only a minor impact on the results.

Allelic ratio (AR) was calculated for each of the informative SNPs (iSNPs) as:

$$AR_{ci} = \frac{\#Alt_{ci}}{\#Alt_{ci}+\#Ref_{ci}}$$

Where c indicates a cell and i indicates an iSNP. #Alt accounts for the number of reads aligned to the non-reference, alternative allele and #Ref accounts for the number of reads aligned to the reference allele. The AR ranges from 0 (only Ref allele) to 1 (only Alt allele).

Each of the iSNPs was assigned by its AR as monoallelic Ref/Alt (AR<=0.1 or AR>0.9 respectively) or biallelic (0.1<AR<=0.9).

For testing the mapping consistency for scRNA-seq data, we compared the number of iSNPs per cell for ChrX and Chr17 (Fig. S2-a). A Pearson correlation of r = 0.68, p-value = 2.23e-15 was calculated for the expression from ChrX and Chr17 (Fig S2-b). The high correlation is an additional confirmation for the reliability of the mapping process. It indicates the coherence in the level of expression associated with each cell. The cumulative fraction of iSNPs (Fig S2-c) indicates that each cell contributes to the overall analysis. Moreover, it emphasized that about 20% of the cells contribute about 50% of the overall cumulative iSNPs in this data set.

##### 1.3.1.4. Identifying escapees in single cell fibroblasts

For genes quantification, we only consider escapees that have multiple evidence for a biallelic expression on ChrX. Specifically, labeling a gene an escapee requires it to have at least 2 biallelic iSNPs over all cells. The same criteria were applied for all autosomes and for Chr17.

The genes, lncRNAs and their chromosomal position were taken from the hg19 UCSC knownCanonical and lincRNAsTranscripts tables (http://hgdownload.cse.ucsc.edu/goldenpath/hg19/database). Note that for the gene centric analysis we removed from our primary hSNP lists those that cover genes on both strands.

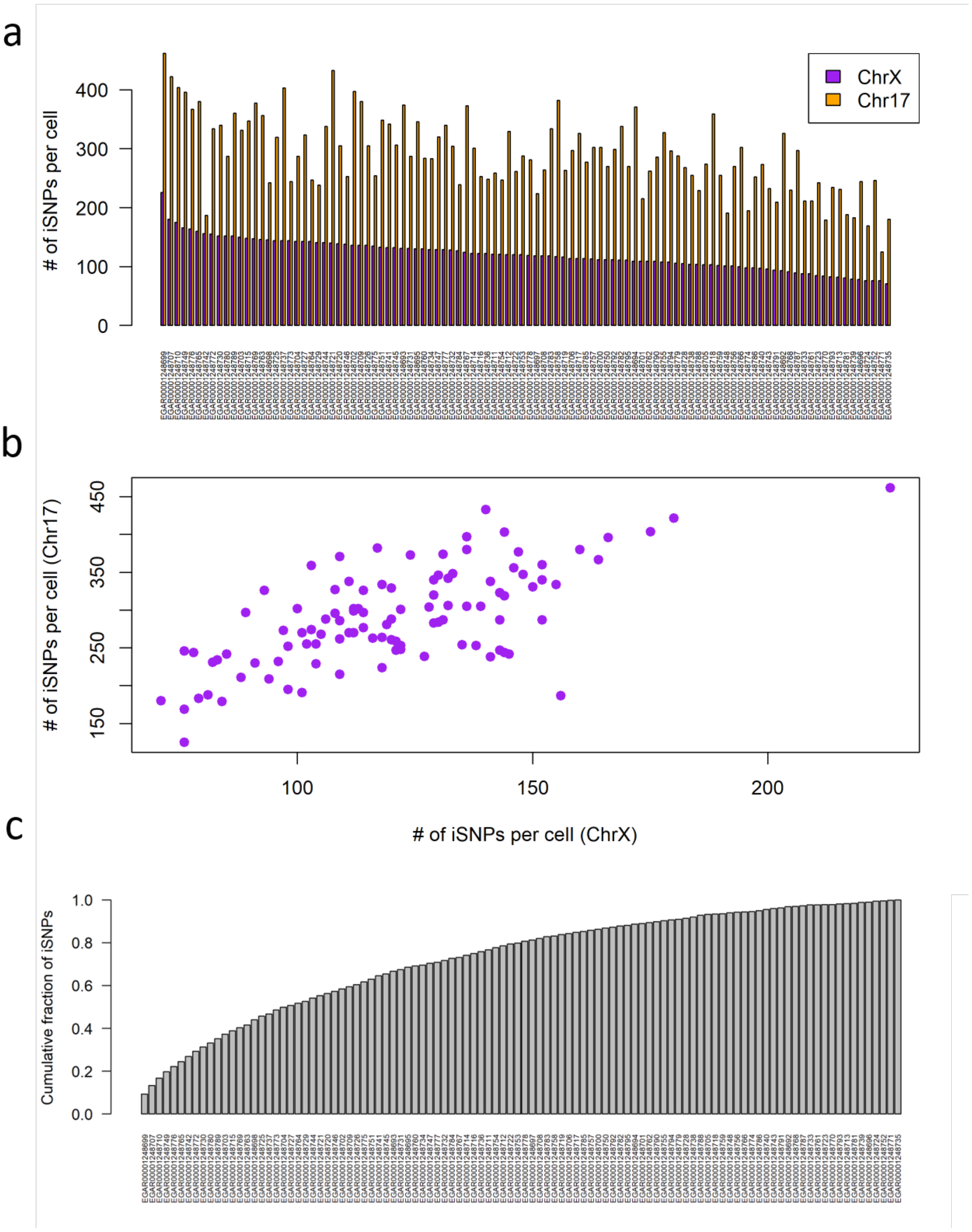


**Fig. S2:** Informative SNPs of 104 single cell primary fibroblasts **(a)** The number of informative SNPs for each of the 104 cells from ChrX and Chr17. Data were collected from 104 fibroblast cells of female origin (UCF_1014). The cells' identifiers are listed in Additional file 2 Supplementary Table S1. **(b)** Correlation between the numbers of SNPs for two chromosomes according to an individual cell. Pearson's correlation of the SNPs for these two chromosomes for all tested cells is r = 0.68, p-value = 2.23e-15. **(c)** Cumulative fraction of iSNPs out of total per each cell. Ordered by the contribution of each of the cells.

##### 1.3.1.5. Cell outliers according to extreme biallelic expression

A possible cause of faulty interpretation for biallelic expression of a gene can originate from a single cell experiment that accidentally included two (or more) cells instead of one cell. Therefore, if for a specific cell, the proportion of biallelic expression shows an extreme value, the data associated from that cell might be explained by having a mixture of two cells expressing two different Xi. We have calculated for each cell the biallelic ratio as *biallelic ratio= biallelic iSNPs/total iSNPs*. The distribution of the cells biallelic expression is indicated in Fig S3. Following this investigation, we have excluded three single cells which had a biallelic ratio of over 3 standard deviations from the average of the entire cells (Fig. S3). After filtering these cells outliers, 101 single cells remained for further analyses.

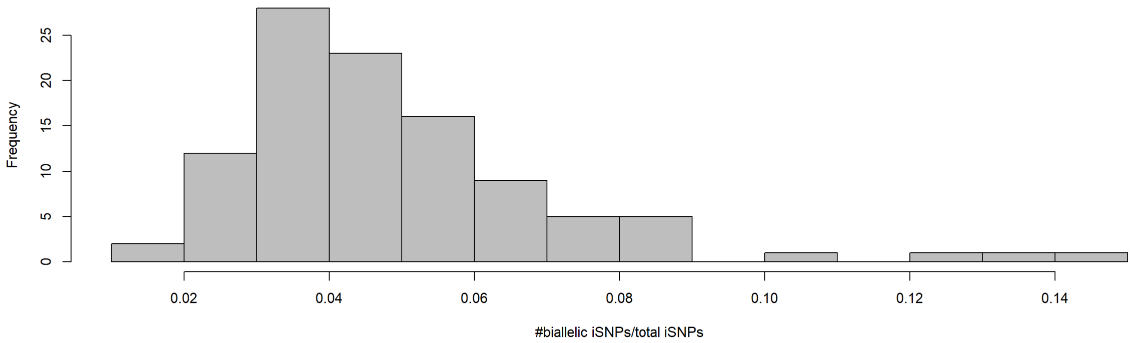


**Fig. S3: The** biallelic ratio of 104 single cell fibroblasts. The histogram shows the cells' frequency of the biallelic ratios. The three most extreme cells on the right were excluded.

#### 1.3.2. Clonal Female Single Cell Lymphoblasts

##### 1.3.2.1. Lymphoblasts and their reference genome

The reference genome used for GM12878 cell line is the diploid NA12878 genome (version Dec 16, 2012, from http://alleleseq.gersteinlab.org/). In short, this genome of the GM12878 cell line is based on hg19, with 4,330,326 SNPs and 829,454 INDELs. The variant list is based on HiSeq 64x sequencing call set from the BROAD institute. Details are available in ftp://gsapubftp-anonymous@ftp.broadinstitute.org/bundle [16, 17]. The diploid genome was extracted from http://sv.gersteinlab.org/NA12878_diploid/NA12878_diploid_genome_2012_dec16.zip.

The allelic phasing is based on the computational pipeline AllelSeq [18]. The paternal and maternal whole haplotypes are used as a reference for the RNA alignment. To achieve the whole genome alignment scheme, the chromosomal sequences were concatenated into a single FASTA file that was indexed by STAR as the reference.

##### 1.3.2.2. Compiling lymphoblast heterozygous SNPs (hSNPs) catalogue

In addition to the diploid genome, we used a VCF file containing all known SNPs for the selected cell line. The file was obtained from http://sv.gersteinlab.org/NA12878_diploid/CEUTrio.HiSeq.WGS.b37.bestPractices.phased.hg19.vcf.gz. From the VCF file, heterozygous loci over all chromosomes were extracted.

Remapping of the VCF coordinates with those of the diploid NA12878 genome (version Dec 16, 2012) was done using LiftOver [6]. Then each SNP was verified as being in that reference position using

Biopython [12] with a window of 5 so that the SNP will be in the center. The remaining SNPs created our list of hSNPs.

The final step for the SNP catalog preparation includes creating a reference GTF file for the positions of the SNPs list on each chromosome. This reference GTF file contains all SNP locations (called GTF_SNP). Formally, in the GTF file, each SNP was considered as having a match with either a paternal feature on the paternal chromosome or a maternal feature on the maternal chromosome.

For the alignment step, chromosomal locations of genes were obtained from UCSC hg19 GTF file. Converting chromosomal locations to parental chromosomal locations was performed using LiftOver available at UCSC toolbox [6]. The conversion created a separate GTF file with maternal and paternal locations of genes. This GTF was then used for STAR alignment.

**1.3.2.3. RNA-Seq realignment and allele counting of single cell clonal lymphoblasts**

Data from 25 single cells RNA-Seq files were downloaded from http://www.ncbi.nlm.nih.gov/geo/query/acc.cgi?acc=GSE44618. RNA-Seq experiments from GM12878 lymphoblastoid cell-line single cells were used [19]. Additional data obtained is a pool of 100 individual cells (Pool100) paired-end RNA-Seq dataset. An identical pipeline was used for the data from Pool100 and single cells after separation of the paired-end reads into single end reads.

For the RNA realignment, FASTQ files were first preprocessed using FASTX (http://hannonlab.cshl.edu/fastx_toolkit/index.html). We have used FASTA_Clipper protocol with the following parameters for trimming out low-score positions (fastq_quality_trimmer -Q33 -t 25 -l 50 –i). SMART adaptors were removed from the sequenced fragments (-Q33 -a AAGCAGTGGTATCAACGCAGAGTACTTTTTTTTT -l 25 -i). The reads were realigned to the paternal and maternal chromosomes using STAR [3] with the default parameters. We eliminated all non-unique mapping (using 'NH:i:1' flag).

**1.3.3.4. Single cell lymphoblast allelic imbalance analysis**

HTSeq pipeline [11] was used for allelic read counting. We used GTF_SNP as a reference for the positions of interest in the genome. HTSeq counted how many of each of the features overlapped the SNPs paternal and maternal locations indicated in the GTF_SNP file. We included identified SNPs from the same read that were labeled as 'ambiguous' (applies for rare instances of closely positioned SNPs). Overall, HTSeq listed for each SNP, the number of aligned reads to maternal or paternal alleles. We activated SamTools for viewing the aligned reads [2].

**1.3.2.5 Allelic imbalance analysis for single cell lymphoblasts**

An Allelic Ratio (AR) was calculated for each iSNP as: $AR_{ci} = \frac{\#Pat_{ci}}{\#Pat_{ci}+\#Mat_{ci}}$

Where c indicates a cell and i indicates an iSNP. #Pat accounts for the number of reads aligned to the paternal allele and #Mat the number of reads aligned to the maternal allele.

We only consider iSNPs that are supported by >=7 reads, aligned at a specific position. Replacing the predetermined read threshold by a cell-specific threshold that was calculated out of the aligned sum of reads for each cell, had only a minor impact on the results.

For visualization purpose, we assigned four types of tags according to the AR associated with each iSNP: (i) paternal; (ii) biallelic expression leaning towards paternal; (iii) biallelic expression leaning towards maternal and (iv) maternal. The tags are set by deciles. Specifically, iSNPs with an allelic ratio of >0.9 and =<0.1 were tagged paternal and maternal, respectively. iSNPs are tagged "biallelic maternal" for 0.1<allelic ratio=<0.5, and as "biallelic paternal" for 0.5<allelic

ratio<=0.9. For some analysis, we combined the tags (ii) and (iii) and iSNPs with an allelic expression ratio of 0.1<ratio<=0.9 are marked 'balanced'.  Identical tagging thresholds were applied for single cells and for a Pool100.

For testing the mapping consistency for scRNA-seq data, we compared the number of iSNPs per cell for ChrX and Chr17 (Fig. S4-a). We observe a Pearson correlation of r = 0.96, p-value = 2.16e-14 between expression from ChrX and Chr17 (Fig. S4-b). Indicating the reliability and coherence of the hSNP mapping between the chromosomes across different cells. Notice that the correlation is higher than for the same analysis that was applied for the fibroblasts (see 1.3.1.3) due to the clonality of the lymphoblasts. The cumulative fraction of iSNPs (Fig. S3-c) indicates that the first 6 cells contribute over 50% of the cumulative information from iSNPs.

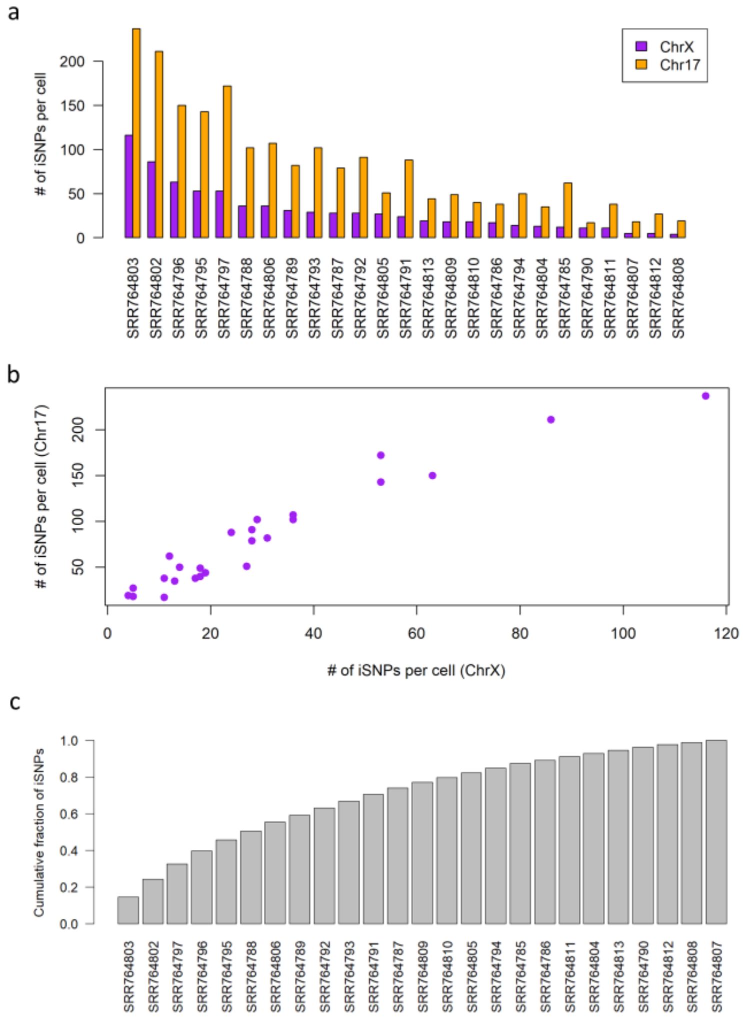


**Fig. S4.** Informative SNPs on ChrX and Chr17 from lymphoblasts **(a)** The number of informative SNPs (iSNPs) for each of the 25 cells. Data were collected from 25 cells of female origin (GM12878 lymphoid cell-line). The cells' identifiers are listed in Additional file 2: Table S1. **(b)** Correlation between the numbers of iSNPs for the two chromosomes according to an individual cell. Pearson's correlation of the

SNPs for these two chromosomes for all tested cells is r = 0.96, p-value = 2.16e-14. **(c)** Cumulative fraction of iSNPs out of total per each cell. Ordered by the iSNP contribution by each cell.

##### 1.3.2.6 Identifying escapees in single cell lymphoblasts

For assigning genes as escapees, we applied two complementary protocols. The first protocol is the strict iSNP protocol that requires multiple iSNPs to be expressed from any of the candidate genes. Multiple iSNPs may represent iSNPs from multiple cells or independent iSNPs in a gene, from the same cell. Due to the sparseness of the hSNPs expressed from single cells, for most instances, multiple iSNPs were collected from different cells. We combine the parental origin allelic evidence into a simplified term called iSNP ratio per gene. This ratio is a simple normalized summation of the iSNPs evidence, where an iSNP tagged as maternal is scored 0 and a paternal/balanced iSNPs are scored 1. The iSNP scores are summed and then normalized by the number of the total iSNPs. We consider escapees as genes with iSNPs score >0 indicating paternal allele expression.

The second protocol is a read-based protocol. For this protocol, the sum of the Paternal and Maternal reads overlapping a gene's hSNPs within its canonical transcript is calculated. A gene will be considered as an escapee if the number of paternal reads is >=7.

The genes, lncRNAs and their chromosomal position were taken from the hg19 UCSC knownCanonical and lincRNAsTranscripts tables (http://hgdownload.cse.ucsc.edu/goldenpath/hg19/database). Note that for the gene centric analysis we removed from our primary hSNP lists those that cover genes on both strands.

#### 1.4. A comparison to a unified annotated ChrX gene catalog

There are 1144 known genes on ChrX [6]. These genes were partitioned into 9 annotated categories [20, 21]. The labels for the 9 categories are: (i) PAR, (ii) escapee, (iii) mostly-escapee, (iv) variable-escapee, (v) mostly-variable-escapee, (vi) conflicting results, (vii) inactivated genes, (viii) mostly-inactivated, and (ix) genes having no data [20, 21]. The annotations are based on a careful analysis according to major publications combining numerous indirect measurements for escapee and inactivated gene identification [22-25]. From all genes in ChrX, 45% have no data, 40% are inactivated-related and the rest carry escapee-associated annotations. We consider escapee-associated collection as a benchmark. This set also includes genes with conflicting evidence and the PAR genes (total 168 genes).

#### 1.5. Statistical analysis

Hypergeometric probability between our results and the external annotated catalog was calculated by comparing the correspondence of any two lists of escapees. We used standard notations of N, k, n and x: N symbolizes all genes on ChrX from [20] with a label other than "No data"; k is the number of escapees by [20] which are associated with any escapee annotations (i.e. the escapee-associated); n is the number of escapees we identified by any of the settings from our protocols; x the number of genes in our list that match the literature-based escapee list in k. P(x) is the probability that an n-trial will result in a value that is > x.

#### 1.6 Additional datasets for annotated escapees

Lists of the annotated escapees according to Tukiainen et al. [26] and Zhang et al [27] were downloaded from the publication supplemental materials. Accordingly, the status of escapee genes was analyzed across the external resources reported [20, 26, 27].
